# From acoustic to linguistic analysis of temporal speech structure: acousto-linguistic transformation during speech perception using speech quilts

**DOI:** 10.1101/589010

**Authors:** Tobias Overath, Joon H. Paik

## Abstract

Speech perception entails the mapping of the acoustic waveform to linguistic representations. For this mapping to succeed, the speech signal needs to be tracked over various temporal windows at high temporal precision in order to decode linguistic units ranging from phonemes (tens of milliseconds) to sentences (seconds). Here, we tested the hypothesis that cortical processing of speech-specific temporal structure is modulated by higher-level linguistic analysis. Using fMRI, we measured BOLD signal changes to 4-s long speech quilts with variable temporal structure (30, 120, 480, 960 ms segment lengths), as well as natural speech, created from a familiar (English) or foreign (Korean) language. We found evidence for the acoustic analysis of temporal speech properties in superior temporal sulcus (STS): the BOLD signal increased as a function of temporal speech structure in both familiar and foreign languages. However, activity in left inferior gyrus (IFG) revealed evidence for linguistic processing of temporal speech properties: the BOLD signal increased as a function of temporal speech structure only in familiar, but not in foreign speech. Network analyses suggested that left IFG modulates processing of speech-specific temporal structure in primary auditory cortex, which in turn sensitizes processing of speech-specific temporal structure in STS. The results thus reveal a network for acousto-linguistic transformation consisting of primary and non-primary auditory cortex, STS, and left IFG.

**Significance Statement:** Where and how the acoustic information contained in complex speech signals is mapped to linguistic information is still not fully explained by current speech/language models. We dissociate acoustic from linguistic analyses of speech by comparing the same acoustic manipulation (varying the extent of temporal speech structure) in two languages (native, foreign). We show that acoustic temporal speech structure is analyzed in superior temporal sulcus (STS), while linguistic information is extracted in left inferior frontal gyrus (IFG). Furthermore, modulation from left IFG enhances sensitivity to temporal speech structure in STS. We propose a model for acousto-linguistic transformation of speech-specific temporal structure in the human brain that can account for these results.

## Introduction

Speech perception entails the mapping of the acoustic waveform to linguistic representations (Poeppel et al., 2008; Kleinschmidt and Jaeger, 2015). Despite decades of research in various disciplines (e.g. linguistics, neuroscience, speech/language development), the mechanisms of this transformation – what aspects of the acoustic speech signal are processed at the neural level, and how they interface with linguistic representations – are still not fully characterized.

The rich temporal structure of speech carries critical information for speech intelligibility (Shannon et al., 1995; Smith et al., 2002), and as such provides a transform from acoustic speech structure to linguistic processes. In speech, linguistic information is conveyed over multiple temporal scales, or temporal windows (Rosen, 1992; Stevens, 2000; Poeppel, 2003; Poeppel et al., 2008). For example, phonemes have an average duration of 30-60 ms, syllables have average durations of 150-300 ms, words are generally longer still, and so forth (Rosen, 1992). These average durations translate to corresponding modulation rates in the speech signal (i.e. ∼17-33 Hz for phonemes, ∼4-7 Hz for syllables, ∼1-3 Hz for words). The slower modulation rates are conveyed via the temporal envelope of the speech waveform and are crucial for speech perception; indeed, if temporal modulations in the delta and theta frequency bands (1-3 and 4-7 Hz, respectively) are removed from the speech envelope, intelligibility breaks down (Ghitza, 2012; Doelling et al., 2014). Conversely, if only temporal envelope modulations are retained, as in noise-vocoded speech (Shannon et al., 1995) or cochlear implants (Wilson, 2004), intelligibility remains robust. Thus, slow modulations, carried in the speech temporal envelope and tracked at the neural level, provide critical information for speech intelligibility.

Recently, we introduced a novel algorithm that controls the temporal extent of natural speech structure via randomizing and then ‘quilting’ back together speech segments of a set duration (Overath et al., 2015). We showed that, whereas earlier processing centers in human auditory cortex – such as Heschl’s gyrus, (HG), part of primary auditory cortex (PAC), or planum temporale (PT), a computational hub receiving information from PAC (Griffiths and Warren, 2002; Kumar et al., 2007; Overath et al., 2007) – are not sensitive to the temporal speech structure, subsequent processing in the superior temporal sulcus (STS) increased as a function of segment length or natural temporal speech structure (Overath et al., 2015). The acoustic manipulation (speech quilting) was performed in a foreign language to participants, and thus addressed neural processes underlying the analysis of acoustic temporal speech structure, irrespective of associated linguistic processes such as syntax, semantics, or lexical access. A dissociation of acoustic versus linguistic processes can be achieved by comparing the same acoustic manipulation in familiar and foreign languages. Controlling both the temporal scale of analysis and the linguistic content in one paradigm ensures that any signal manipulations will affect acoustic properties of the speech signal similarly in both languages; in contrast, such signal manipulations will affect linguistic processes only in the familiar language.

To date, only few studies have taken this approach. For example, listeners are able to track hierarchical linguistic structure based on syntax and semantics only in a familiar, but not in a foreign language (Ding et al., 2015); this is also reflected in oscillatory entrainment to natural familiar speech, which is strongest in the delta band (Pérez et al., 2015). However, since most of these studies only investigated effects of language familiarity in either continuous speech (Peña and Melloni, 2012; Ding et al., 2015; Pérez et al., 2015) or intact single words (Strelnikov et al., 2011), they were unable to reveal which temporal scales in the speech signal are critical for, and amenable to, linguistic modulation.

Here we utilize speech quilts in familiar and foreign languages to map how processing speech-specific temporal structure proceeds from acoustic analysis to linguistic analysis. Dissociating acoustic and linguistic processes in this manner reveals which temporal scales (e.g. phonemic, syllabic, words) in the speech signal interface with linguistic representations, and how the latter modulates the former.

## Methods

### Participants

21 participants (mean age = 23.57, range = 19-28, 9 females) were native speakers of American English, with no knowledge of Korean. All reported to have normal hearing and no history of neurological or psychiatric diseases. Two participants were excluded from further analysis: one participant performed at chance for the speaker identification task in the scanner, while the other only completed 3 runs, leaving a total of 19 participants (mean age = 23.68, range = 19-28, 9 females). Participants provided written consent prior to participating in the study in accordance with the Duke University Health System Institutional Review Board.

### Stimuli

Sounds were derived from recordings (44100 Hz sampling rate, 16 bit resolution) of four bilingual female English/Korean speakers reading from a book in either language (native English and Korean speakers judged the recordings as coming from native speakers). This removes voice cues for language identification. The recordings were then used as source material for the quilting algorithm (Overath et al., 2015). Briefly, a source signal is divided into equal-length segments, which are then pseudorandomly rearranged, or stitched together, to create a new speech quilt signal. By using an L^2^ norm when choosing adjacent segments to approximate the original segment-to-segment change in the original speech signal, and by using pitch-synchronous overlap-add (PSOLA(Moulines and Charpentier, 1990)) to avoid sudden frequency jumps at segment boundaries, the quilting algorithm ensures that low-level acoustic attributes (e.g. amplitude modulation rate, frequency spectrum) in the speech quilt are similar to those in the original speech signal. For both languages (English and Korean), the stimuli of the 5 experimental conditions are 4 s long speech quilts made up of 30 ms, 120 ms, 480 ms, or 960 ms speech segments, as well as 4 s long original, unaltered excerpts from the recordings. The choice of segment lengths sub-samples those used in (Overath et al., 2015), while also allowing a confirmation of the response plateau at ∼500 ms for foreign speech, and testing its validity for speech-specific processing in a familiar language.

### Experimental design

Prior to the main experiment in the scanner, participants were familiarized with the four speakers in a behavioral experiment. Trials consisted of original, unaltered 4 s long recordings of the four speakers and were presented via Sennheiser HD 380 pro headphones using Psychophysics Toolbox (Brainard, 1997) in Matlab. During the first couple runs, participants saw the speaker identity while they listened to each trial (e.g. ‘Speaker 1’) on the monitor. During subsequent runs, participants identified speaker identity for each trial via pressing keys 1-4 on the keyboard without any visual cues; feedback was provided for each trial (correct, wrong). Each individual run took approximately 4 minutes, and participants needed to reach at least 90% correct performance to proceed to the fMRI experiment.

The 10 experimental conditions from a 2 Language (English, Korean) x 5 Segment length (30 ms, 120 ms, 480 ms, 960 ms, Original) factorial design were presented in eight “runs”, each lasting ∼6.5 minutes. Stimuli were presented in a pseudo-randomized fashion that boosted contrast selectivity, e.g. by ensuring that presentations of the 30 ms segment length speech quilt and original speech conditions of each language were close together in time (not more than two trials apart; contrasts between trials that are far apart in time can potentially be affected by the high-pass filter, see below). Each experimental condition was presented 32 times per scanning session (in addition to 32 silent trials of 4 sec duration) with a mean inter-stimulus interval of 4 s (randomly sampled from a uniform distribution between 3-5 s). All stimuli were unique and were presented only once. Stimuli inside the scanner were presented using Psychophysics Toolbox(Brainard, 1997) running in Matlab at a comfortable listening level (∼75 dB SPL) at 44100 Hz sampling rate and 16 bit resolution via Sensimetrics (www.sens.com) MRI-compatible insert earphones (Model S14); participants wore protective earmuffs to further reduce the background noise of the scanner environment.

In the scanner, participants’ again performed the speaker identity task by pressing one of four buttons on an MRI-compatible button box to indicate which speaker they had heard for each trial; while participants had been trained on the original, unaltered 4 s long recordings, stimuli in the scanner also included the speech quilt stimuli. Participants were instructed to only register their response after the sound had ended (to avoid confounding the BOLD signal response with a motor execution response), and were given feedback on each trial (correct, wrong, missed) and for each run (percentage correct).

### Image acquisition

Data were recorded on a GE MR750 3.0 Tesla scanner using an 8-channel head coil and a high-resolution echo-planar imaging (EPI) sequence yielding contiguous isotropic 2×2×2 mm voxels (110×110 matrix, FOV = 22, TE = 28 ms, flip angle = 90°, TR = 2.2s). 36 slices were acquired for each volume in an interleaved ascending sequence to avoid signal bleeding between adjacent slices. The volume was centered on STG and spanned from the inferior colliculus (IC) to inferior frontal gyrus (IFG). A high-resolution 1×1×1 mm voxel-size T1-weighted MRI (FSPGR) scan (TR/TE: 2,089/3.18 ms, FOV: 256) was acquired for each participant to inform structure-function mapping.

### Data analysis

Imaging data were analyzed using Statistical Parametric Mapping software (SPM12, http://www.fil.ion.ucl.ac.uk/spm). The first four of the 174 volumes in each run were discarded to control for T1 saturation effects. The remaining 1360 scans were realigned to the first volume in the first run, un-warped to correct for motion artifacts and re-sliced using sinc interpolation (SPM12, “Realign and Unwarp”), and slice time corrected to account for differences in slice acquisition time (SPM12, “Slice timing”); the structural scan of each participant was coregistered to the mean functional scan (SPM12, “Coregister”) and segmented into grey and white matter and cerebro-spinal fluid and spatially normalized to standardized stereotaxic MNI space (SPM12, “Segment”), before applying the resulting linear transformations to the EPIs and structural scan (SPM12, “Normalize: Write”). Finally, the EPIs were spatially smoothed to improve the signal-to-noise ratio using an isotropic 6 mm full-width at half-maximum (FWHM) Gaussian kernel (SPM12, “Smooth”).

The design matrix for each participant consisted of 10 regressors (corresponding to the 10 experimental conditions), which were derived by convolving the stimulus (modeled as a four-second box-car function) with SPM’s canonical hemodynamic response function. The silent periods were not modeled explicitly. Data were high-pass filtered at 1/256 Hz to remove slow drifts in the signal.

Standard second-level group analyses in SPM were based on a random-effects model within the context of the general linear model(Friston et al., 1995). For second-level group analyses, the smoothing of first-level functional contrast images was increased to an effective 8 mm FWHM Gaussian kernel to better allow for inter-individual differences in anatomy.

In addition, we calculated the BOLD signal in a set of anatomically and functionally defined ROIs using MarsBaR (Brett et al., 2002). Two cortical anatomical ROIs in HG (encompassing primary auditory cortex) and PT (part of non-primary auditory cortex) were based on published probability maps in Rademacher et al. (2001) and Westbury et al. (1999), respectively. Both ROIs were thresholded such that they only included voxels with at least 30% probability of belonging to either structure. Two subcortical anatomical ROIs in early auditory structures in IC and the medial geniculate body (MGB) were spherical ROIs (with a radius of 5 mm) centered on published coordinates of these structures ([−6 −34 −12] and [6 −34 −12] for IC (Griffiths et al., 2001); [−16 −28 −8] and [16 −28 −8] for MGB (Devlin et al., 2006)). For ROI analyses in these subcortical structures, the data were not smoothed.

For each participant, the BOLD signal in each run to the 5 conditions of each language (English, Korean) was normalized to the mean BOLD signal of their original, unaltered condition in the other 7 runs. For example, for the English speech conditions, the BOLD signal for Eng30, Eng120, Eng480, Eng960, EngOrig in run 8 was normalized to the mean BOLD signal for EngOrig in runs 1-7. This was then averaged across runs for each participant.

To investigate the response in the functional ROI (fROI) in STS, we employed a leave-one-out procedure to avoid double-dipping (Kriegeskorte et al., 2009): we defined separate fROIs for English and Korean conditions via [EngOrig > Eng 30ms] and [KorOrig > Kor 30 ms] functional contrasts, respectively, in 7 runs, leaving out the data from the eighth remaining run; this was done 8 times (once for each left-out run). Only voxels that a) survived a significance threshold of p < 0.001 (uncorrected for multiple comparisons across the volume) and b) lay within the superior temporal lobe were evaluated. The response in each left-out run and for each language was then normalized with respect to the mean response to the EngOrig or KorOrig condition in the 7 runs that formed the fROI, respectively. For example, for the English speech conditions, the BOLD signal for Eng30, Eng120, Eng480, Eng960, EngOrig in run 8 was normalized with respect to the mean BOLD signal for EngOrig in runs 1-7. This procedure was repeated for the other 7 leave-one-out permutations. The results were then averaged across runs for each participant.

The procedure was similar for the fROI in left IFG: for the [EngOrig > Eng 30 ms] functional contrast for 7 runs, only voxels in the left-out run that a) survived a significance threshold of p < 0.005 (uncorrected) and b) lay within a mask defined by BA44 and BA45 (from the Anatomy Toolbox, version 1.5 (Eickhoff et al., 2005)) were included in the analysis. For some participants who had no supra-threshold voxels in a given 7-run fROI (each participant had 8 possible fROIs), we randomly chose one voxel within the mask; this was the case for 3 participants in 1, 3, and 1 runs, respectively. This procedure ensures that statistics can be run, while simultaneously penalizing data from those participants.

For the psychophysiological interaction (PPI) analysis (Friston et al., 1997), we chose as seed the fROI defined in the left IFG for each participant, as described above. We then searched for areas that were modulated by activity in left IFG as a function of segment length.

### Statistics

Normalized BOLD percentage signal change data in ROIs, and behavioral accuracy (percentage correct) data were analyzed via two-way repeated-measures (RM) ANOVAs, using the Greenhouse-Geisser correction in cases where Mauchly’s test indicated significant violations of the assumption of sphericity.

## Results

### Acoustic analysis (effects of temporal speech structure)

We first searched for areas that showed an increase in activity as a function of increasing temporal speech structure. Figure 1 shows this for the [EngOrig > Eng30ms] and [KorOrig > Kor30ms] group functional contrasts and reveals areas in STS for both English and Korean, as well as left IFG for English speech. To enable a direct comparison with the previous study (Overath et al., 2015), which did not include original, natural speech, we also investigated the [Eng960ms > Eng30m] and [Kor960ms > Kor30ms] group functional contrasts; the pattern of results was very similar to that revealed in Figure 1.

**Figure 1:**
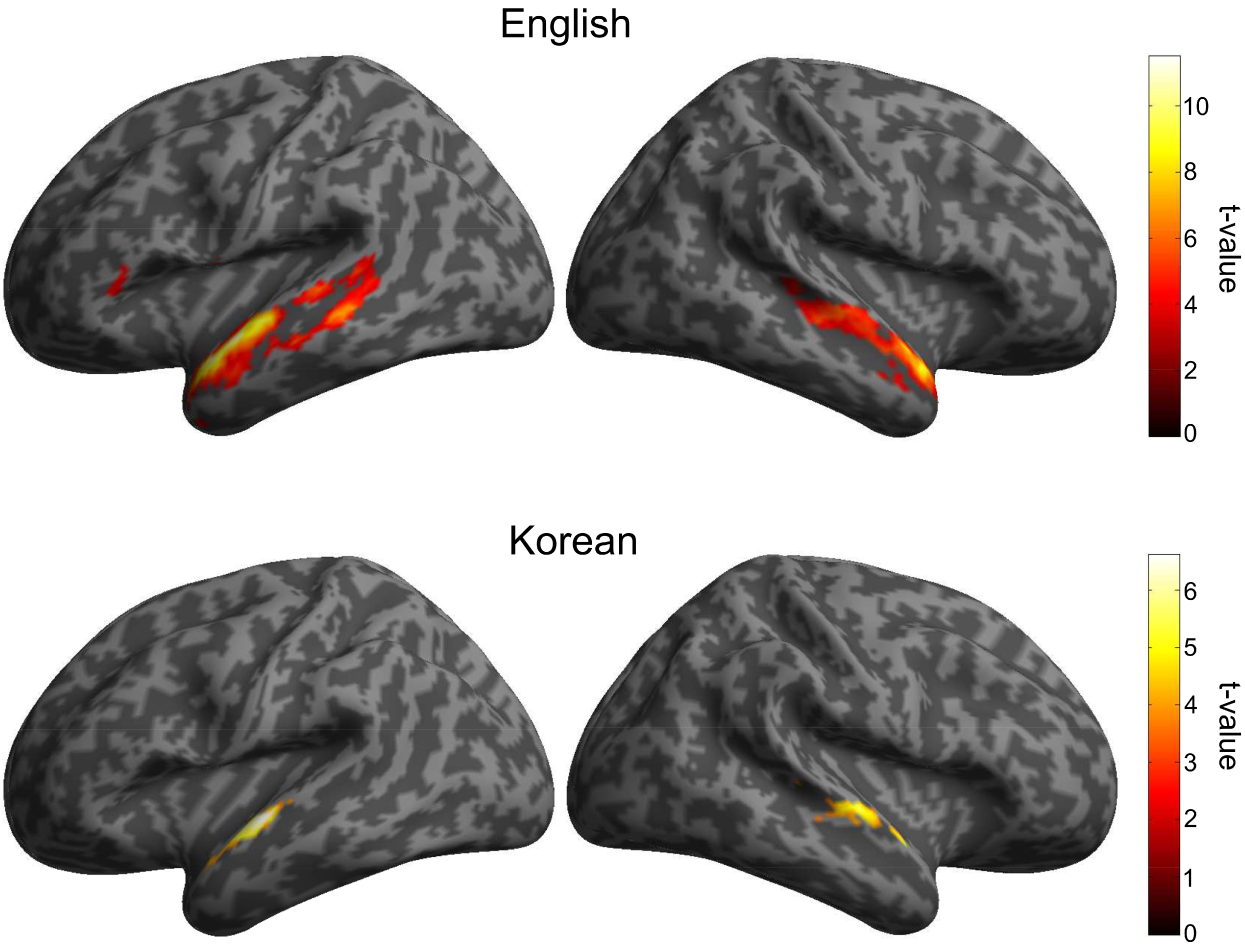
Areas showing significantly stronger BOLD signal to original speech compared to speech quilted with 30 ms segment lengths in English (top) and Korean (bottom).

Next, we investigated the response in these fROIs located along STS, as well as in anatomically defined cortical ROIs of the auditory system, i.e. HG and PT. The responses in cortical ROIs and fROIs differed significantly (main effect of ROI: F_(2.08,37.43)_ = 173.45, p < 0.001, η^2^_p_ = 0.91), and we therefore investigated their responses separately. The BOLD signal in ROIs of early cortical auditory areas (HG and PT) decreased slightly as a function of segment length (Figure 2): RM ANOVAs for HG and PT with factors Hemisphere, Segment length, and Language revealed weak main effects of Segment length (F_(4,72)_ = 4.01, p = 0.005, η^2^_p_ = 0.18; F_(2.59,46.67)_ = 4.68, p = 0.008, η^2^_p_ = 0.21; for HG and PT, respectively); for Korean only, post-hoc pairwise comparisons revealed a significant difference (p < 0.05, Bonferroni corrected) in PT, and only between the 30 ms and 480 ms speech quilt conditions, and between the 30 ms and original speech conditions.

**Figure 2:**
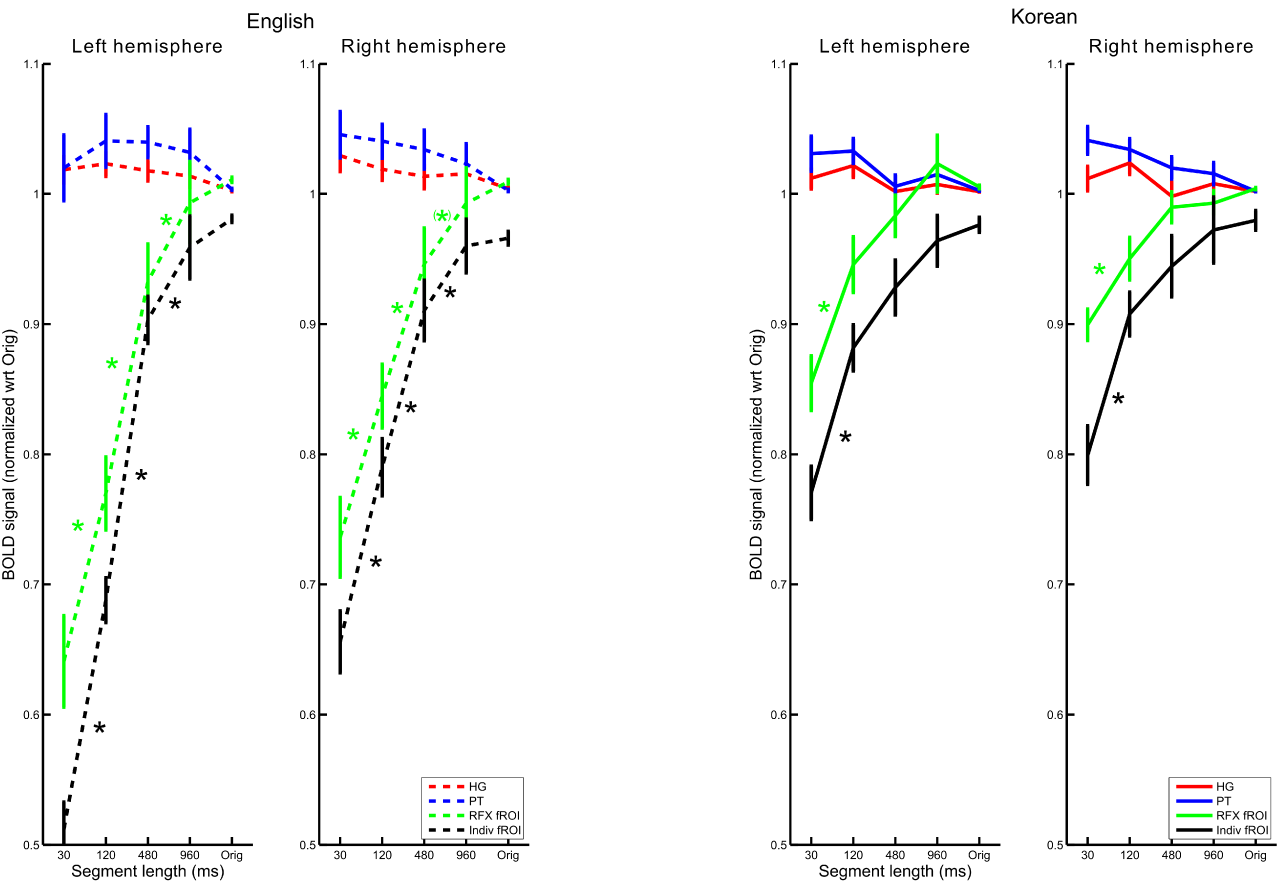
Response in anatomical ROIs HG (red) and PT (blue), and functional ROIs of the group fROI shown in Figure 1 (green) and individual fROIs (black), shown separately for the two languages and two hemispheres. Error bars denote ±1 SEM. Asterisks denote significant pairwise comparisons (after Bonferroni correction), p < 0.05. For each language, responses are normalized within each ROI to the response to original speech in the left-out run (see Methods).

In the group fROI for English, the BOLD signal increased as a function of temporal speech structure (main effect of Segment length: F_(4,72)_ = 62.57, p < 0.001, η^2^_p_ = 0.78) and differed between hemispheres (main effect of Hemisphere: F_(1,18)_ = 7.37, p = 0.01, η^2^_p_ = 0.29); the effect of Segment length was more pronounced in the left hemisphere (interaction: F_(2.22,40.03)_ = 9.52, p < 0.001, η^2^_p_ = 0.35). In the group fROI for Korean, the BOLD signal increased as a function of Segment length (F_(2.45,44.09)_ = 20.49, p < 0.001, η^2^_p_ = 0.53), while revealing an interaction with Hemisphere (F_(4,72)_ = 6.16, p < 0.001, η^2^_p_ = 0.26).

For the individual fROIs for English and Korean the pattern was largely identical, but the size of the effects was generally larger. For the individual English fROI, the BOLD signal increased as a function of segment length (F_(2.87,51.7)_ = 163.49, p < 0.001, η^2^_p_ = 0.9), differed between hemispheres (F_(1,18)_ = 12.37, p = 0.002, η^2^_p_ = 0.41), and revealed an interaction (F_(4,72)_ = 25.71, p < 0.001, η^2^_p_ = 0.59). Post-hoc pairwise comparisons between adjacent segment length conditions showed significant differences (p < 0.05, Bonferroni corrected) between all but the 960 ms and original speech conditions.

For the individual Korean fROI, the BOLD signal increased as a function of segment length (F_(4,72)_ = 48.04, p < 0.001, η^2^_p_ = 0.73). Post-hoc pairwise comparisons between adjacent segment length conditions showed significant differences (p < 0.05, Bonferroni corrected) on the left between 30 ms, 120 ms, and 480 ms conditions, and on the right between 30 ms and 120 ms conditions.

The coverage of fMRI volume also allowed us to investigate the sensitivity to temporal speech structure in earlier, subcortical structures of the auditory system (this was not possible in Overath et al. (2015)). Figure 3 shows that IC and MGB were unaffected by the manipulation of speech structure in the speech quilts.

**Figure 3:**
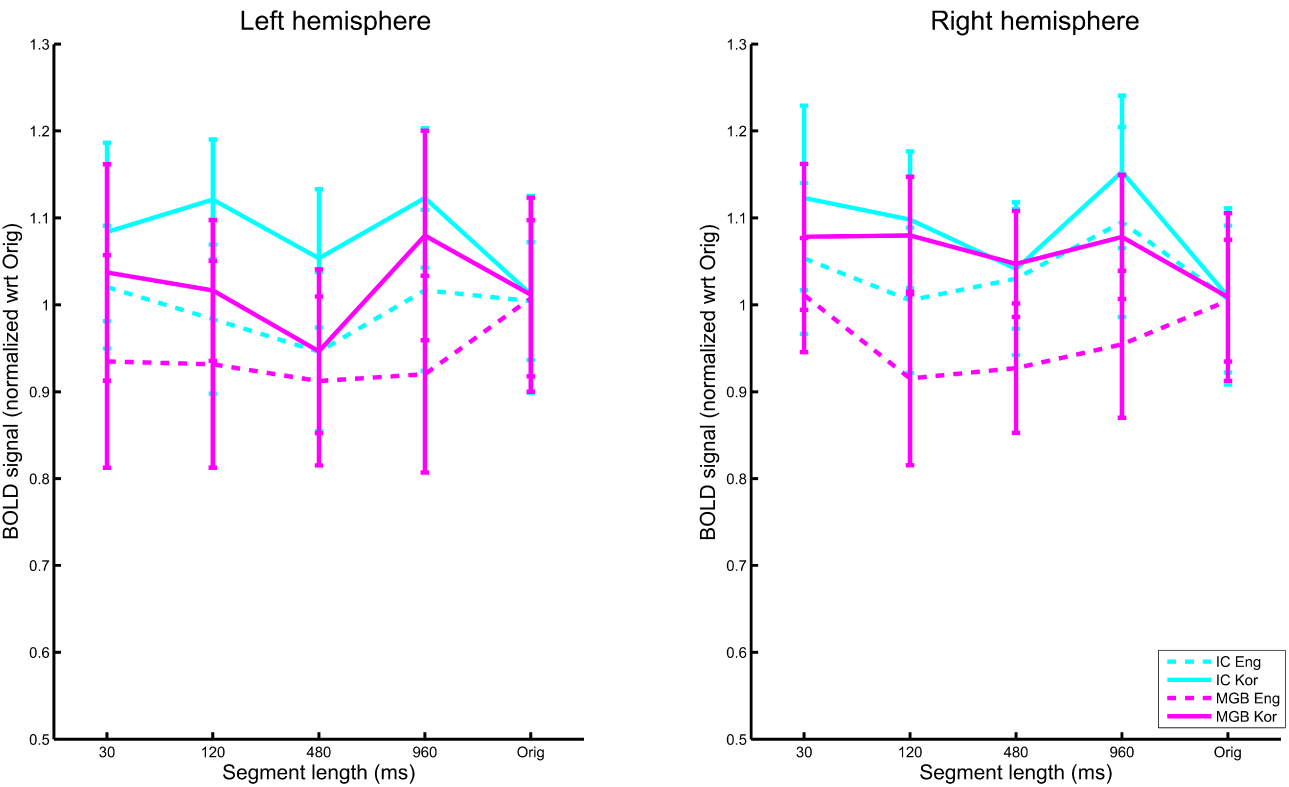
Response in IC and MGB. Error bars denote ±1 SEM. For each language, responses are normalized within each ROI to the response to original speech in the left-out run (see Methods).

### Linguistic analysis (effects of language familiarity)

The results presented so far concern effects of temporal speech structure that are similar in nature for English and Korean; that is, they address an analysis at the level of signal acoustics, irrespective of language familiarity. Next, we searched for neural responses that differ between languages.

The response in the fROIs revealed three notable differences between languages: First, the size of the effect of segment length was larger for English than Korean (main effect of ROI: F_(1,18)_ = 64.14, p < 0.001, η^2^_p_ = 0.78). Second, the response in the individual STS fROI for Korean plateaued at ∼480 ms (in fact, it seemed to decrease for original Korean, though this was not statistically significant), replicating the earlier finding of Overath et al. (2015) for a foreign language; however, for English the response only plateaued at twice the segment length, 960 ms. Third, the volume of the individual fROI in STS was significantly larger in the left hemisphere than in the right only for in English: a RM ANOVA revealed main effects of Language (F_(1,18)_ = 42.37, p < 0.001, η^2^_p_ = 0.7) and Hemisphere (F_(1.67,29.99)_ = 15.27, p < 0.001, η^2^_p_ = 0.46), as well as an interaction (F_(2.55,46)_ = 16.74, p < 0.001, η^2^_p_ = 0.48).

Beyond the response in STS, the functional contrast [EngOrig > Eng30ms] also revealed an effect of segment length in the left IFG (see Figure 1): the response in left IFG increased as a function of segment length only for English, but was flat for Korean (the [KorOrig > Kor30ms] functional contrast did not reveal any effects beyond the temporal lobes) (Figure 4). A RM ANOVA revealed main effects of Language (F_(1,18)_ = 171.31, p < 0.001, η^2^_p_ = 0.91) and Segment length (F_(4,72)_ = 38.2, p < 0.001, η^2^_p_ = 0.68), which was due to a significant interaction (F_(4,72)_ = 28.76, p < 0.001, η^2^_p_ = 0.62). Post-hoc pairwise comparisons between adjacent conditions revealed that, for English, only 30 ms vs. 120 ms, and 480 ms vs. 960 ms were not significantly different (p > 0.05, Bonferroni corrected). Furthermore, between languages, all comparisons except for speech quilts with 30 ms segment length were significant (p < 0.05, Bonferroni corrected).

**Figure 4:**
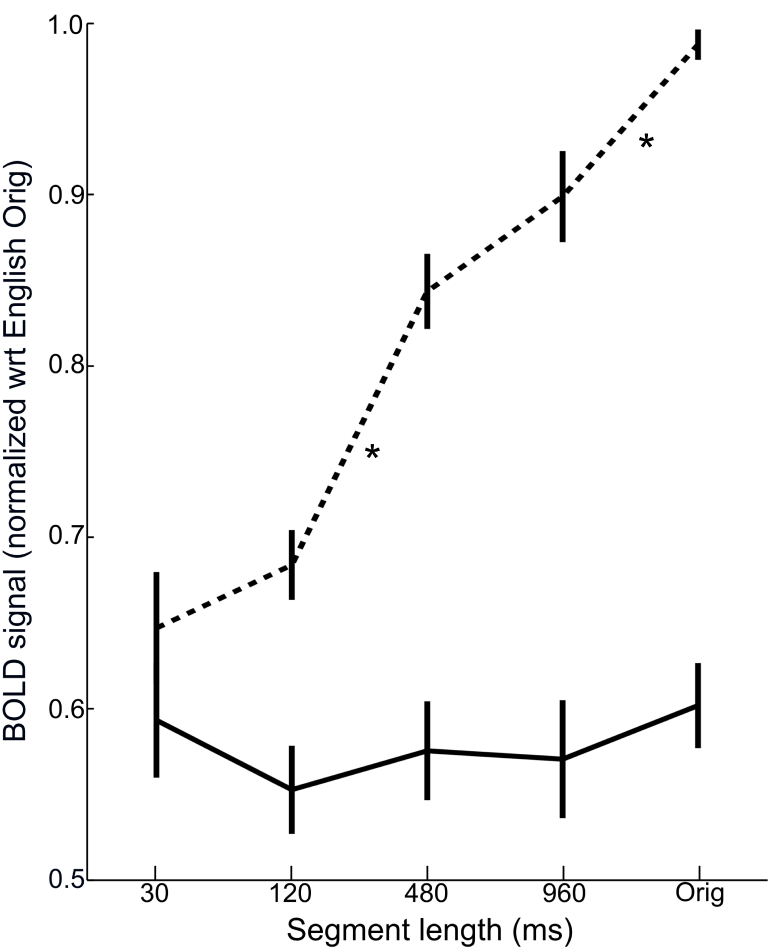
Response in individually defined (leave-one-out) fROIs in left IFG to speech quilts and original speech in English (dotted) and Korean (solid). Error bars denote ±1 SEM. Asterisks denote significant pairwise comparisons (after Bonferroni correction), p < 0.05, within language.

### Functional Connectivity

We investigated whether the response in left IFG modulated the response in auditory areas in the temporal lobe via a PPI analysis (Friston et al., 1997). This revealed that modulation of primary and non-primary auditory cortex (bilateral HG, PT, and STG) by left IFG increased as a function of segment length (Figure 5). Note that the areas in auditory cortex that display a modulatory effect by segment length are largely distinct from those revealed by the English group fROI (yellow) and show only small overlap (white).

**Figure 5:**
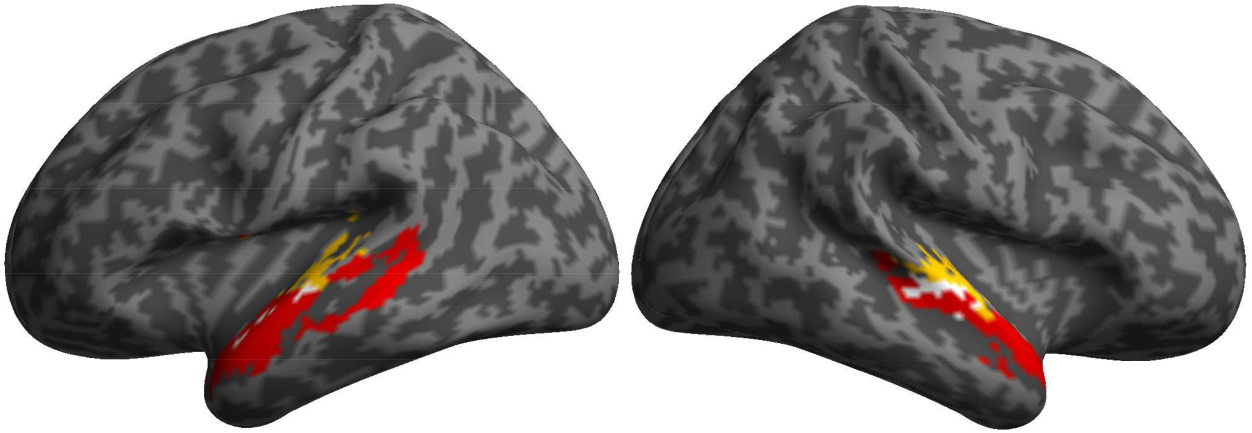
Regions showing significant modulation by segment length (in English) (yellow) and a stronger response to English original speech than English speech quilted with 30 ms segments (red); areas of overlap are shown in white.

### Effects of task

We also investigated areas that showed a stronger response to short temporal speech structure than original speech via the functional contrast [30ms > Orig] (the [Eng30ms > EngOrig] and [Kor30ms > KorOrig] functional contrasts did not differ significantly from each other). This revealed a bilateral network of areas in the middle and anterior insula, inferior parietal cortex, and a small region in PT (Figure 6).

**Figure 6:**
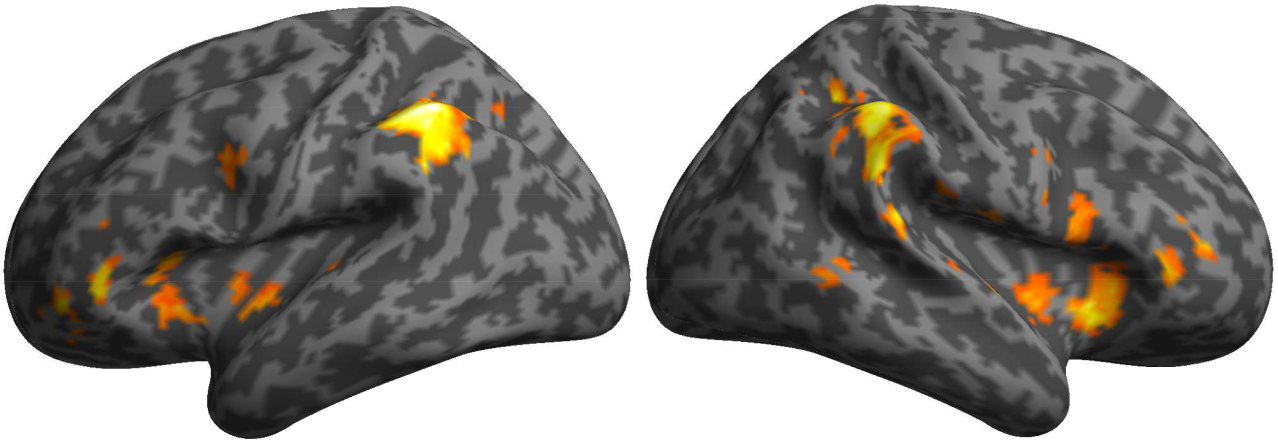
Areas showing significantly stronger BOLD signal to speech quilted with 30 ms segment lengths compared to original speech (across languages).

### Behavioral results

Average behavioral performance in the speaker identification task (mean: 81.82% correct performance, std: 14.38%) was well above chance (25%). Performance generally increased with segment length, which was reflected in a main effect of Segment length (RM ANOVA: F_(4,72)_ = 10.23, p < 0.001, η^2^_p_ = 0.36), and was better for English than Korean (main effect of Language: F_(1,18)_ = 35.79, p < 0.001, η^2^_p_ = 0.67). Performance for Korean speech quilts was slightly more variable, leading to a weak Language x Segment length interaction (F_(4,72)_ = 2.62, p = 0.04, η^2^_p_ = 0.13).

## Discussion

We provide evidence for the neural processes underlying the transformation from acoustic analysis to linguistic analysis of temporal speech structure. The results reveal that STS processes the acoustic properties of temporal speech structure, while left IFG maps this acoustic information to linguistic representations. In addition, connectivity analyses suggest that this mapping modulates the processing of acoustic speech properties in earlier auditory areas in cortex.

Based on decades of research, current speech/language models are now able to delineate the major cortical structures and their putative roles in speech perception and production (Hickok and Poeppel, 2007; Peelle, 2012; Skeide and Friederici, 2016). However, while there is some evidence for how they interact and modulate each other, e.g. as a function of intelligibility (Leff et al., 2008; Park et al., 2015; Tuennerhoff and Noppeney, 2016), it is currently still unclear how they proceed from the analysis of speech-specific acoustic structure to linguistic analysis (acoustic-linguistic transformation). This is mainly because the majority of studies on which current speech/language models are based either only manipulated linguistic content (e.g. via syntactic or semantic violations) or both linguistic and acoustic content (e.g. via noise vocoding, spectral rotation, time-reversed speech).

In contrast, the current study was able to dissociate acoustic from linguistic analyses of temporal speech structure by applying the same acoustic manipulation in familiar and foreign languages. Thus, simultaneously controlling (1) the temporal scale at which analysis occurs (via speech quilting) and (2) the linguistic content (via two different languages), ensured that neural responses that vary as a function of segment length, but are shared or similar for the two languages, would suggest an analysis at the signal-acoustics level; however, neural responses that differ based on language familiarity would imply the presence of linguistic processing.

The results confirm that acoustic analysis of temporal speech structure takes place in STS: the BOLD signal increased as a function of temporal speech structure in both foreign and familiar languages. In general, the effects in STS for the current foreign language (Korean) were somewhat weaker than for the foreign language (German) in Overath et al. (2015). For example, while previously the normalized BOLD signal for speech quilts with 30 ms segment lengths was about 60% of that for 960 ms segment lengths, it was about 80% in the current study. Similarly, the current study revealed a significant difference between segment lengths only for the two shortest segment lengths used. These differences may be related to a number of factors. First, the effect estimation in Overath et al. (2015) was more robust due to a total of 52 scanning sessions (and up to 4 scanning sessions of a given participant), as opposed to 19 scanning sessions in the current study. Second, the etymological difference between Korean and English is greater than that between German and English, and it is possible that the corresponding differences in temporal speech acoustics between Korean and English resulted in reduced neural engagement. In fact, we chose Korean precisely because of its etymological distance to English, so as to obtain as ‘clean’ a measure of acoustic analysis of temporal speech structure as possible. Future studies will therefore have to explore further the degree to which the etymological dis/similarity between languages affects the acoustic processing of speech structure in STS.

Overath et al. (2015) demonstrated that the BOLD signal plateaued for segment lengths greater than ∼500 ms. Importantly, since the inflection point was the same for time-compressed speech quilts – which, in a given segment, contain twice as much temporal structure as normal uncompressed speech – the plateau was attributed to intrinsic analysis properties of auditory cortex, rather than reflecting the analysis of intrinsic stimulus properties. In the current study, the BOLD signal did not show a clear plateau and generally continued to increase beyond 480 ms segment lengths. However, it should be noted that, while this trend was visible for Korean, it did not reach statistical significance. In contrast, the response to 960 ms speech quilts in English was significantly larger than that to 480 ms speech quilts, in both the left and right hemispheres.

It is possible that the inclusion of the speaker identification task, and the associated increase in task difficulty and attention, may have led to a generally enhanced sensitivity to longer temporal windows of analysis. Future studies will need to determine whether analysis windows beyond ∼500 ms are indeed malleable to task demands or attention, for example via a direct comparison of attended vs. ignored speech quilts as a function of segment length.

Similarly, a novel finding in the current study concerns the role of a task on the response in auditory cortex and beyond. The results in Overath et al. (2015) did not reveal any areas that showed a stronger response as a function of decreasing segment length. Whereas that study simply asked participants to press a button at the end of each sound, the behavioral speaker identification task in the current study required participants to engage with the stimulus more explicitly (while still being orthogonal to the acoustic manipulation of temporal speech structure and language familiarity). In fact, speaker identification became more challenging and performance decreased as the segment length decreased. The areas that showed a stronger response to speech quilts with short segment lengths are associated with processing demands in linguistic tasks (Falkenberg et al., 2011; Yue et al., 2013). Thus, it is possible this effect is less due to the analysis of temporal acoustics in speech signals, but more related to general attentional or task demands.

While the response increase in STS as a function of segment length was bilateral for foreign speech (see also Overath et al., 2015), its extent was significantly left-lateralized for familiar speech. The (pre-linguistic) acoustic analysis of temporal speech structure thus takes place in both hemispheres; however, if linguistic processes are able to become engaged, then left-hemispheric structures in STS are more strongly recruited. This provides further evidence for the view that speech perception is largely a bilateral process, for which lateralization emerges only once linguistic processes become engaged (Peelle, 2012).

The limited temporal resolution of fMRI provides a summary measure of the cortical response to temporal speech structure; however, it cannot speak to the underlying neural mechanisms that track the temporal speech signal with millisecond precision. This will require more direct measurements of neural activity that provide high temporal resolution (e.g. M/EEG, ECoG). A potential neural mechanism underlying the parametric BOLD signal increase in STS is that speech envelope entrainment increases as a function of temporal speech structure. In particular, the temporal envelope of natural speech contains prominent amplitude modulations in the delta (1-3 Hz) and theta (4-7 Hz) frequency bands that carry important linguistic units, roughly corresponding to words and syllables, respectively (Greenberg et al., 2003; Goswami and Leong, 2013; Ding et al., 2017). Recent evidence suggests that such neural entrainment to the temporal speech envelope correlates with speech intelligibility (Ding and Simon, 2014; Di Liberto et al., 2015; Vanthornhout et al., 2018). Thus, future studies will need to investigate to what degree neural entrainment is dependent on acoustic vs. linguistic processes.

Based on the current results we propose a model for acousto-linguistic transformation of speech-specific temporal structure in the human brain in which auditory information is passed from primary and non-primary auditory cortices to STS for processing of temporal speech structure; this stage of processing is primarily concerned with the analysis of acoustic properties of temporal speech structure. If, however, this information contains familiar linguistic information (e.g. phonemic, lexical, semantic, syntactic cues), it is passed on to left IFG for linguistic processing. Left IFG in turn modulates the processing in earlier auditory areas (such as HG, PT, and STG), which in turn induces greater sensitivity in STS for speech-specific temporal structure; this would explain the steeper slope for increasing speech-specific temporal structure for familiar speech (English) compared to foreign speech (Korean).

There is recent evidence in support of this view, both in terms of a temporal progression of linguistic analysis from temporal to left frontal cortices, and top-down modulation of auditory cortex. For example, the analysis of acoustic speech information in temporal cortex precedes phonological analysis in left inferior frontal cortex (Toscano et al., 2018), while top-down signals from frontal cortex increase the ability of auditory cortex to track temporal speech envelope modulations in the delta and theta bands (Park et al., 2015).

In conclusion, by simultaneously controlling temporal speech structure and linguistic familiarity, the current study was able to disambiguate the neural contributions underlying acoustic and linguistic processing of temporal speech structure. The results thereby inform our understanding of where and how linguistic information interfaces with, and modulates the temporal analysis of speech.

## Acknowledgements

This work was supported in part by US National Institutes of Health grant 1R21DC016386 to T.O.The authors thank Kimberly Paige Mihalsky for assistance with data collection.

## Notes

The authors declare no financial or other conflicts of interest.

